# Structural consequences of BMPR2 kinase domain mutations causing pulmonary arterial hypertension

**DOI:** 10.1101/786756

**Authors:** Apirat Chaikuad, Chancievan Thangaratnarajah, Frank von Delft, Alex N Bullock

**Author notes:** **Corresponding author**. Alex N Bullock,.

## Abstract

Bone morphogenetic proteins (BMPs) are secreted ligands of the transforming growth factor-β (TGF-β) family that control embryonic patterning, as well as tissue development and homeostasis. Loss of function mutations in the type II BMP receptor BMPR2 are the leading cause of pulmonary arterial hypertension (PAH), a rare disease of vascular occlusion and heart hypertrophy. To understand the structural consequences of these mutations, we determined the crystal structure of the human BMPR2 kinase domain at 2.35 Å resolution. The structure revealed an activate conformation of the catalytic domain that formed canonical interactions with the bound ligand Mg-ADP. Disease-associated missense mutations were mapped throughout the protein structure, but clustered predominantly in the larger kinase C-lobe. Modelling revealed that the mutations will destabilize the protein structure by varying extents consistent with their previously reported functional heterogeneity. The most severe mutations introduced steric clashes in the hydrophobic protein core, whereas those found on the protein surface were less destabilizing and potentially most favorable for therapeutic rescue strategies currently under clinical investigation.

## Introduction

Pulmonary arterial hypertension (PAH) is a devastating cardiovascular disease that affects up to 15 patients per million population ^1^. The pathology typically presents in individuals aged 30-50 years who suffer vascular occlusion and high pressure in the pulmonary arteries due to the abnormal proliferation of pulmonary artery endothelial cells, pulmonary artery smooth muscle cells and fibroblasts ^2^. Most patients succumb to heart hypertrophy and cardiac failure within 3-5 years of diagnosis.

To date, sequencing studies have identified 16 causative genes for PAH ^3^. Mutations in the gene *BMPR2* have emerged as the predominant risk factor accounting for some 53–86% of heritable cases and 14–35% of idiopathic cases ^4^. The majority of these involve autosomal dominant non-sense and frameshift mutations that implicate *BMPR2* haploinsufficiency as the pathogenic mechanism ^3^. *BMPR2* encodes for the bone morphogenetic protein (BMP) type II receptor kinase (BMPR2), which assembles with type I BMP receptor kinases to transduce BMP ligand signaling through the phosphorylation of transcription factors SMAD1/5/8 ^5^. Significantly, loss of function mutations associated with PAH have also been identified in the type I BMP receptor ALK1 (*ACVRL1*), their co-receptor endoglin (*ENG)* and their physiological ligands BMP9 (*GDF2*) and BMP10 (*BMP10*) ^3^. These factors assemble specifically in endothelial cells to form a functional heteromeric signaling complex suggesting the loss of BMP9/10 signaling as the basis for vascular dysfunction ^6^.

Over 400 different PAH-associated mutations have now been identified in the *BMPR2* gene ^7^. Importantly, some 25% of these are missense mutations where a protein product is likely to be expressed without non-sense mediated decay. These mutations are distributed throughout the gene affecting both the extracellular ligand-binding domain and the intracellular kinase domain.

Functional studies have revealed dominant negative behavior, as well as heterogeneity in the effects of mutation ^8–10^. Cysteine substitutions disrupting the extracellular cystine knot domain were found to cause retention of mutant BMPR2 within the endoplasmic reticulum, whereas kinase domain mutants were properly trafficked to the plasma membrane but lacked functional signaling through the SMAD1/5/8 pathway. An interesting exception was the kinase domain mutant E503D, which activated a SMAD-dependent transcriptional reporter construct as efficiently as wild-type BMPR2 ^10^.

To better understand the structural consequences of PAH-associated missense mutations we determined the crystal structure of the BMPR2 kinase domain at 2.35 Å resolution. The structure revealed three classes of mutations, including those causing buried steric clashes, those removing buried packing interactions and those found on the protein surface. Modelling of the mutants indicated that the surface mutations were the least destabilizing and therefore the most favorable for potential therapeutic rescue strategies under current clinical investigation^11^.

## Results

### Structure Determination

To enable structural studies, the kinase domain of human BMPR2 was cloned into a bacterial expression vector providing a non-cleavable C-terminal hexahistidine tag and purified by Ni-affinity, size-exclusion and anion exchange chromatography. Diffracting crystals were obtained in the presence of the ligand ADP and the crystal structure was solved and refined at a resolution of 2.35 Å. Two protein molecules were present in the asymmetric unit. Both chains formed an identical interaction with Mg-ADP, but showed minor differences in the partially disordered activation loop as well as the C-terminus through which crystal contacts were formed. A summary of statistics for data collection and refinement is reported in Table 1.

**Table 1.**
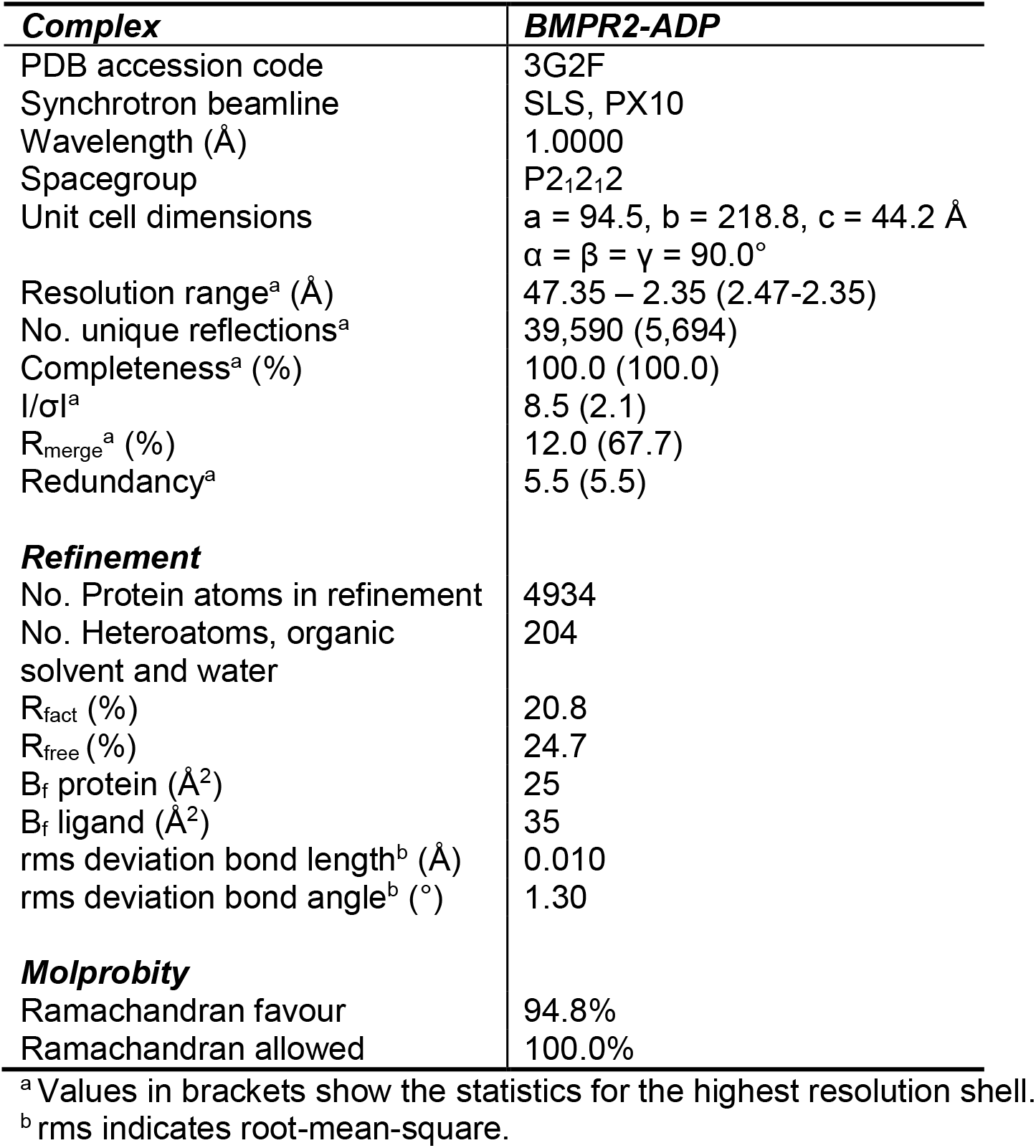
Data collection and refinement statistics

### Conserved features of the BMPR2 kinase domain

BMPR2 shows the typical bilobal architecture of a protein kinase. The structure reveals a conserved fold with other type II receptors with a pattern of specific kinase domain insertions that define the BMP/TGF-β receptor family (Figure 1). These include the L45 loop and the E6 loop as well as insertions flanking the αF helix and an insertion in the substrate pocket preceding the αG helix.

**Figure 1.**
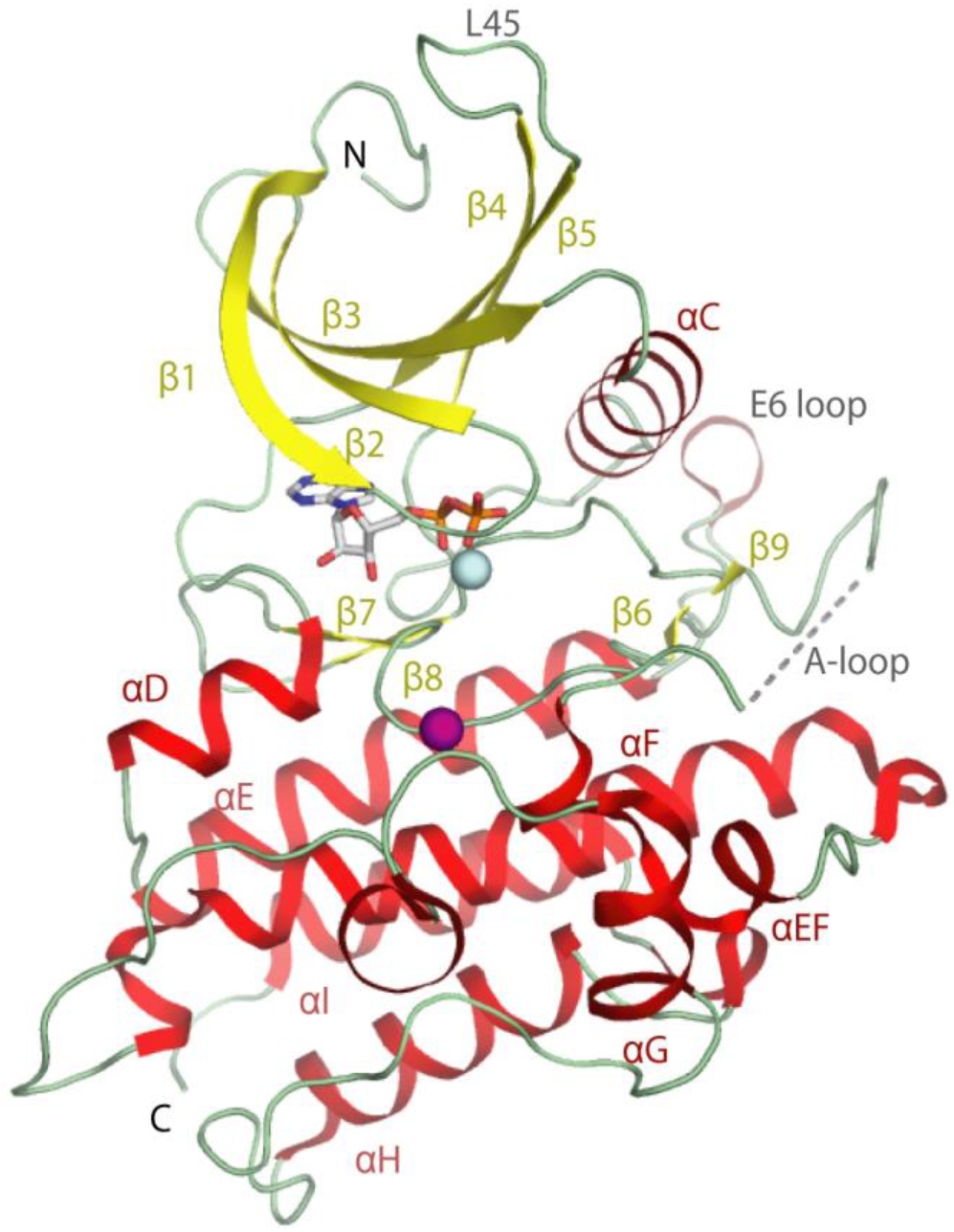
Structural overview of the BMPR2-ADP complex. The secondary structure elements of BMPR2 are labelled and shown as ribbons. ADP is shown in stick representation, and Mg^2+^ ion (cyan) and sulphate molecule (purple) are displayed as spheres. Disordered residues in the activation loop are indicated by a dotted grey line.

### The BMPR2 kinase domain adopts an active conformation

The type II receptor BMPR2 lacks the regulatory GS domain found in BMP type I receptors and is accepted to be constitutively active ^5^. In agreement, the BMPR2 structure displays features characteristic for an active kinase (Figure 2). The N and C-terminal kinase lobes adopt a closed conformation. The correct positioning of the αC helix is evidenced by the 2.8 Å salt bridge formed between the catalytic residues Lys230 (β3 strand) and Glu243 (αC helix) (Figure 2). Canonical interactions are also observed with Mg-ADP, including two hydrogen bonds with the kinase hinge region and further hydrogen bonds with the catalytic loop and phosphate-binding loop (β1-β2), also known as the glycine-rich loop (Figure 3). The activation loop (A-loop) makes surprisingly few contacts with the catalytic domain. Here, interaction with the αC helix is mediated by a hydrogen bond formed between Asn242 and the carbonyl of Ser355 (Figure 2). The BMP/TGF-β receptor kinases differ from many other kinase families in that they are not regulated by activation loop phosphorylation and therefore lack the common interaction between this phosphate moiety and the αC. For this reason, we predict that further interaction with substrate is necessary to order this A-loop segment. Of note for such interaction, the A-loop in BMPR2 contains a six residue insertion relative to activin type II receptors, such as ACVR2B (Figure 2). By contrast, the loop conformation is well defined in the equivalent ACVR2B structure ^12^ and appears stiffened by three proline residues that are not conserved in BMPR2 (Figure 2).

**Figure 2.**
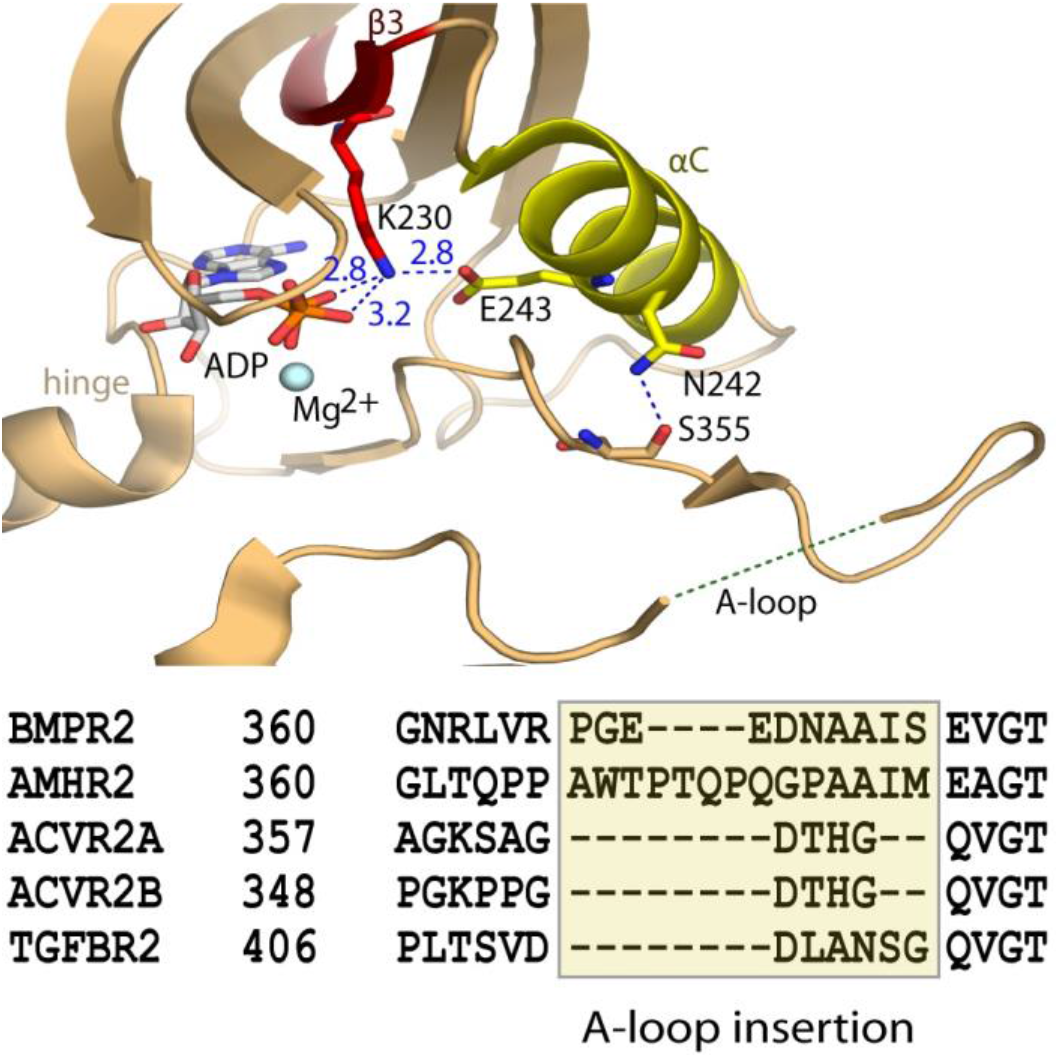
Active conformation of the BMPR2 catalytic domain. (A) BMPR2 adopts an active conformation as shown by a salt bridge between the catalytic residues K230 and E243. Electrostatic hydrogen bond and salt bridge interactions are denoted by a blue dashed line together with their distances (Å). A large insertion in the activation loop is disordered. A sequence alignment reveals insertions here in the activation loops of BMPR2 and AMHR2 compared to other type II receptors.

**Figure 3.**
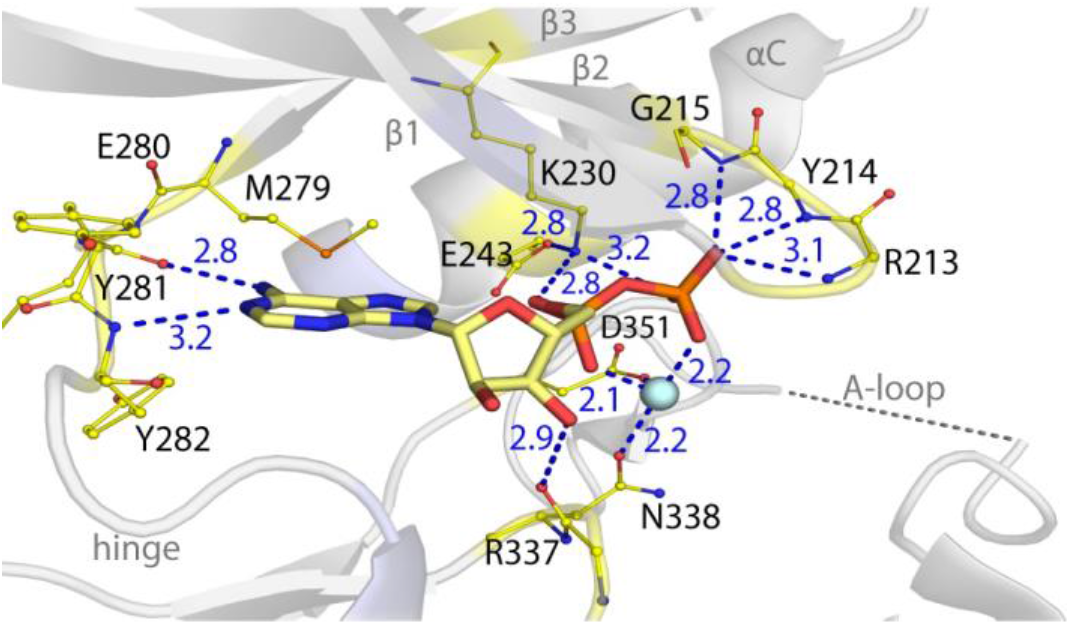
ATP-pocket interactions of the bound Mg-ADP. BMPR2 forms canonical interactions with ADP and a single Mg^2+^ ion (cyan sphere). These include two hydrogen bonds with the kinase hinge region as well as further hydrogen bonds with the catalytic loop and phosphate-binding loop (β1-β2).

### PAH-associated mutations destabilize the active BMPR2 structure

PAH-associated missense and nonsense mutation sites are located throughout the length of the BMPR2 protein. Due to their large number, we focus our analysis here on the subset of missense mutations reported by Machado et al. that fall within the kinase domain of BMPR2 (Figure 4) ^13^. The structure of BMPR2 reveals how these PAH-associated mutations effect a loss of function.

**Figure 4.**
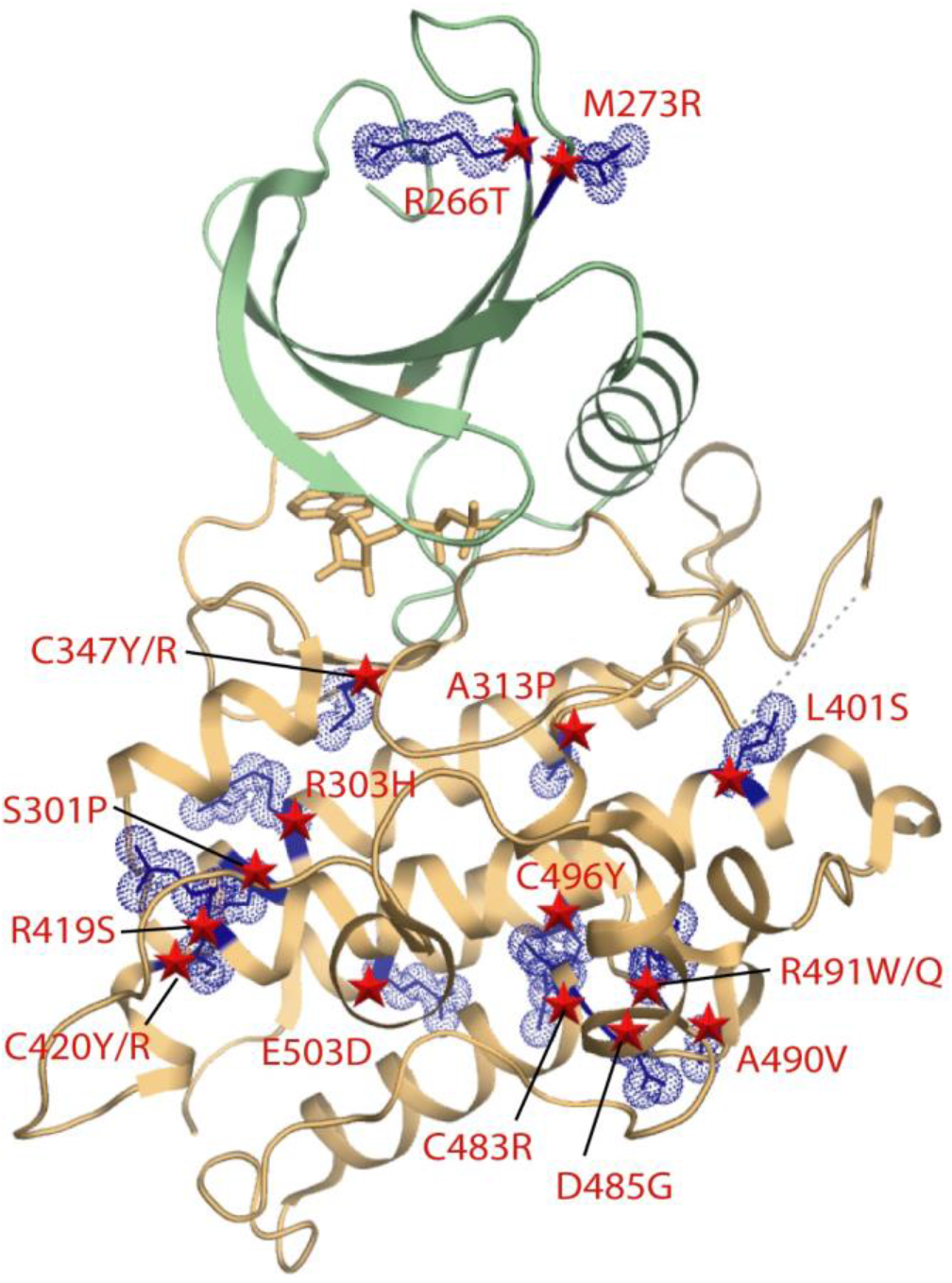
Locations of PAH-associated missense mutations in the BMPR2 structure. Ribbon representation is shown for the kinase N and C-lobes and coloured green and brown, respectively. The positions of mutations are indicated by blue stick and space-fill representations of the wild-type side chains. The distribution of PAH-associated mutations in BMPR2 is widespread in the kinase C-lobe, but more limited in the N-lobe.

The most frequent missense mutations are buried in the kinase C-lobe. These mutations are predicted to severely disrupt the stability and folding of the catalytic domain (Table 2). Here, the most destabilizing mutations S301P, A313P, C347R/Y, C420R/Y, C483R and R491W are all predicted to introduce severe steric clashes (Figure 5). Ser301 is located near the start of the long αE helix where it forms backbone and side chain hydrogen bonds with the backbone carbonyl of Asp297 to stabilize both the helix and the preceding loop conformation. Its mutation to proline would break both of these interactions, introduce local steric clashes and truncate the αE helix. Ala313 lies in the same αE where it packs against the C-terminal αI helix. A mutation to proline at this position would similarly truncate the αE helix resulting in the perturbation of the adjacent ATP-binding pocket. Notably, four of the most severe mutations affect cysteine residues in the hydrophobic core of the kinase C-lobe. The C347R and C347Y mutations located within the β8 strand introduce large bulky side chains that would form steric clashes with residues in the surrounding β7 strand, as well as with helices αD, αE and αF. Similar bulky substitutions arising from the C420R and C420Y mutations would affect the end of the αF helix and are predicted to disrupt proper folding through severe steric clashes with Trp298 (αE). The fourth cysteine position Cys483 (αH) packs in the hydrophobic core against residues in the αE and αI helices. Its mutation to arginine would introduce unsatisfied charge into the core and cause a severe clash with Tyr413 (αF). The final severe clash is introduced by the R491W mutation, which occurs within the loop connecting helices αH and αI. This mutation would disrupt a salt bridge with Glu386 in the short αEF helix, as well as hydrogen bonds to the backbone carbonyl of Asp485, another site of disease mutation.

**Figure 5.**
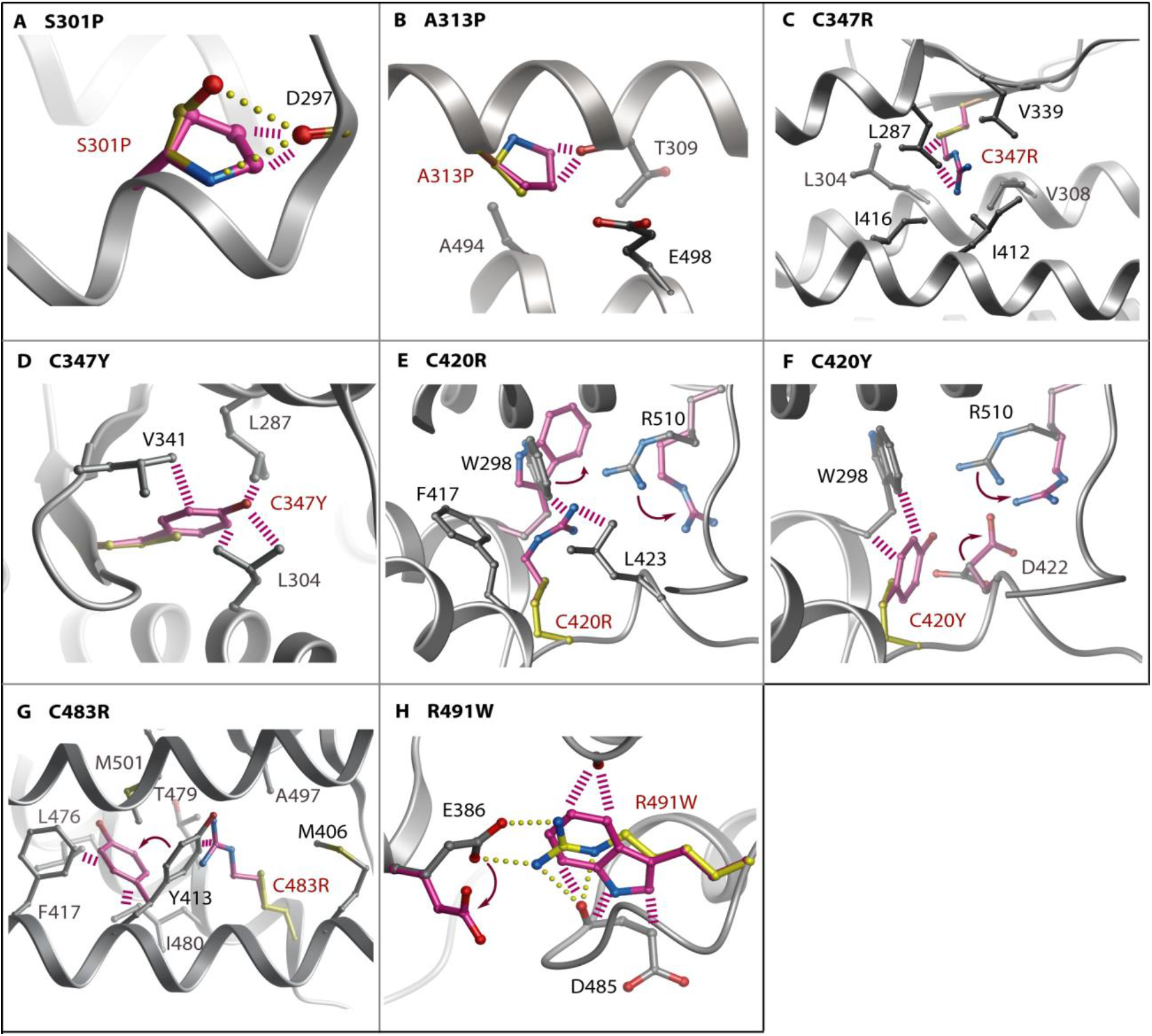
Modelling of PAH-associated mutations causing buried steric clashes. Structural models were calculated using the Eris server (http://dokhlab.unc.edu/tools/eris/) ^25^. Changes in the mutant structure are coloured purple and overlaid onto the wild-type residue coloured yellow. Hydrogen bonds in the wild-type (yellow) and clashes in the mutant structure (purple) are shown by spheres and dashed lines, respectively.

**Table 2.** Predicted destabilization of PAH-associated mutants of BMPR2

| **Class** | **Mutation site** | **Predicted $\Delta\Delta G^a$** |
| --- | --- | --- |
| **1. Buried steric clashes** | S301P | N.D.<sup>b</sup> |
|  | A313P | >10 |
|  | C347R | >10 |
|  | C347Y | >10 |
|  | C420R | >10 |
|  | C420Y | >10 |
|  | C483R | >10 |
|  | R491W | >10 |
| **2. Buried broken interactions** | L401S | 4.97 |
|  | R419S | 7.41 |
|  | R491Q | 5.94 |
| **3. Exposed** | R266T | 0.78 |
|  | M273R | 2.08 |
|  | R303H | 5.57 |
|  | D485G | 7.97 |
|  | A490V | 5.56 |
|  | C496Y | 4.93 |
|  | E503D | 1.66 |

<sup>a</sup> Structural models and destabilization free energies were calculated using the Eris server (http://dokhlab.unc.edu/tools/eris/). The $\Delta\Delta G$ value defines the difference in the free energy between the wild-type and mutant structures. Values >0 are destabilizing.
<sup>b</sup> Not determined. The S301P mutation was too severe for the Eris server to model. The structural model was instead calculated using ICM <sup>26</sup>.

A further three PAH-associated mutations L401S, R419S and R491Q also affect buried sites in the protein structure, but result in smaller side chain substitutions that are predicted to disrupt packing interactions without introducing steric clashes (Figure 5 and Table 2). Leu401 is located near the start of the large αF helix where it contributes to hydrophobic packing with the β9 strand. Its mutation to serine would break these van der Waals contacts and potentially perturb the proximal catalytic loop by forming a new hydrogen bond to Arg332. This arginine position falls within the highly conserved HRD motif of the eukaryotic kinases, which functions in catalysis by engaging the bound substrate as well as the kinase activation loop. Arg419 packs at the end of the αF helix where it forms 3 hydrogen bonds to tether the large αG-αH loop in close proximity to the shorter αD helix. Introduction of a serine at this position would break these bonds and disrupt the folding around this part of the kinase C-lobe. The R491Q mutation is predicted to be slightly less damaging than the R491W mutation described above, but will nonetheless break the salt bridge with Glu386 (αEF helix) to disrupt folding.

The remaining mutation sites are solvent exposed and generally predicted to be less disruptive (Figure 5 and Table 2). A notable exception is the mutation D485G which breaks two hydrogen bonds that help to order the αH-αI loop. Perturbation of this loop risks disrupting two further backbone hydrogen bonds to R491, another site of disease mutation described above. R266T and M273R are more unusual in that they are located in the smaller N-lobe of the kinase domain. Arg266 (β4) forms hydrogen bonds with backbone carbonyls in the β1 strand that would be lost upon mutation causing some destabilization of the N-lobe β-sheet conformation. Met273 is an exposed residue in the β5 strand. Mutation to arginine introduces potential hydrogen bonding to Glu265 (β4) which may cause minor perturbation of the surface packing. The R303H (αE), A490V (αH-αI loop), C496Y (αI) and E503D (αI) mutations are not predicted to affect hydrogen bonding. Rather these substitutions may subtly influence the side chain packing at the surface of the kinase domain. The occurrence of apparently benign mutations at the protein surface suggests a potential role for some of these residues in protein-protein interactions. Further functional and structural studies are needed to define these interaction partners and their binding sites.

## Discussion

Here we present a molecular model for the BMPR2 receptor catalytic domain and its deregulation in disease. BMP/TGF-β signaling is unusual in utilizing two distinct classes of transmembrane receptor kinases, the type II and type I receptors, respectively, which form a heteromeric complex for ligand binding and intracellular signaling. Biochemical assays have suggested that type II receptors such as BMPR2 are constitutively active, whereas the type I receptors require activating phosphorylations within their juxtamembrane GS domains by the type II receptors. Consistent with this view, the kinase domain structure of BMPR2 shows an active conformation as evidenced by the correct positioning of the αC helix, as well as the catalytic residues within the ATP-binding pocket which are captured in complex with Mg-ADP. However, a minor deviation from this state is apparent from some localized disorder in the activation loop. This observation potentially reflects the requirement for this flexible region to bind to the substrate type I receptor as they receptors assemble together upon extracellular ligand binding. Additional structures will be needed to elucidate the precise nature of this most important interaction.

PAH-associated mutations have been shown to abrogate receptor trafficking, assembly and downstream SMAD signaling ^8–10^. The structure reveals how many of the missense mutations break interactions in the core of the BMPR2 structure to disrupt protein folding. Importantly for our understanding of the genotype-phenotype relationship, the structure also explains the contrasting effects of a subset of solvent-exposed mutations. Thus, the functionally defective mutant D485G disrupts a site of four critical hydrogen bonds, while the mutation E503D has no apparent structural consequences and has proven functional in cellular SMAD reporter assays ^10^. This class of mutation may potentially interfere with BMPR2 function in alternative ways, for example disrupting other protein-protein interactions, as observed for C-terminal tail mutants which fail to bind to the kinase LIMK1 ^14^.

Overall, all of the analyzed mutations were predicted to be destabilizing to some degree consistent with the accepted disease model of BMPR2 loss of function. Current rescue strategies under investigation include the use of small molecule chaperones to maximize receptor trafficking, as well the delivery of recombinant BMP9 to bolster the remaining signaling potential ^6,11^. The varying extent of the predicted disruption suggests that different mutations will have different propensities for rescue. In this respect, we identify a subset of surface exposed mutations that would appear to be the most promising for further investigation.

## Methods

### Protein expression and purification

The kinase domain of human BMPR2 (residues 189-517) was subcloned into the pNIC-CH vector and transformed into *E. coli* strain BL21(DE3)R3-pRARE2 for expression. Cultures were induced with 1 mM IPTG overnight at 18°C and the cells harvested and lysed by ultrasonication. Recombinant proteins were purified by Ni-affinity, size-exclusion and anion exchange chromatography. Proteins were stored at 4°C buffered in 50 mM Hepes pH 7.4, 300 mM NaCl, 10% glycerol, 10 mM DTT, 50 mM L-arginine, 50 mM L-glutamate.

### Crystallization

Crystallization of the BMPR2-ADP complex was achieved at 4°C using the sitting-drop vapour diffusion method. BMPR2 was pre-incubated with 10 mM ADP at a protein concentration of 5.8 mg/mL, and crystallized using a precipitant containing 25% PEG 8,000, 0.3 M ammonium sulphate, 0.05 M MgCl_2_, and 0.1 M sodium cacodylate pH 6.0. Diffraction quality crystals grew in a 150 nL crystallization drop containing an equal volume of the protein and reservoir solution.

### Data collection and structure refinement

Crystals were cryoprotected with mother liquor plus 20% ethylene glycol for the BMPR2-ADP complex and vitrified in liquid nitrogen. Diffraction data were collected at Swiss Light Source, station PX10 using monochromatic radiation at wavelength 1.000 Å. Data were processed with MOSFLM ^15^ and subsequently scaled using the program SCALA from the CCP4 suite ^16^. Initial phases were obtained by molecular replacement using the program PHASER ^17^ and the structure of ACVR2B (PDB 2qlu) as a search model. Density modification and NCS averaging were performed using the program DM ^18^, and the improved phases were used in automated model building with the programs ARP/wARP ^19^ and Buccaneer ^20^. The resulting structure solution was refined using REFMAC5 from the CCP4 suite ^21^ and manually rebuilt with COOT ^22^. Appropriate TLS restrained refinement using the tls tensor files calculated from the program TLSMD ^23^ was applied at the final round of refinement. The complete structure was verified for geometric correctness with MolProbity ^24^.

**Figure 6.**
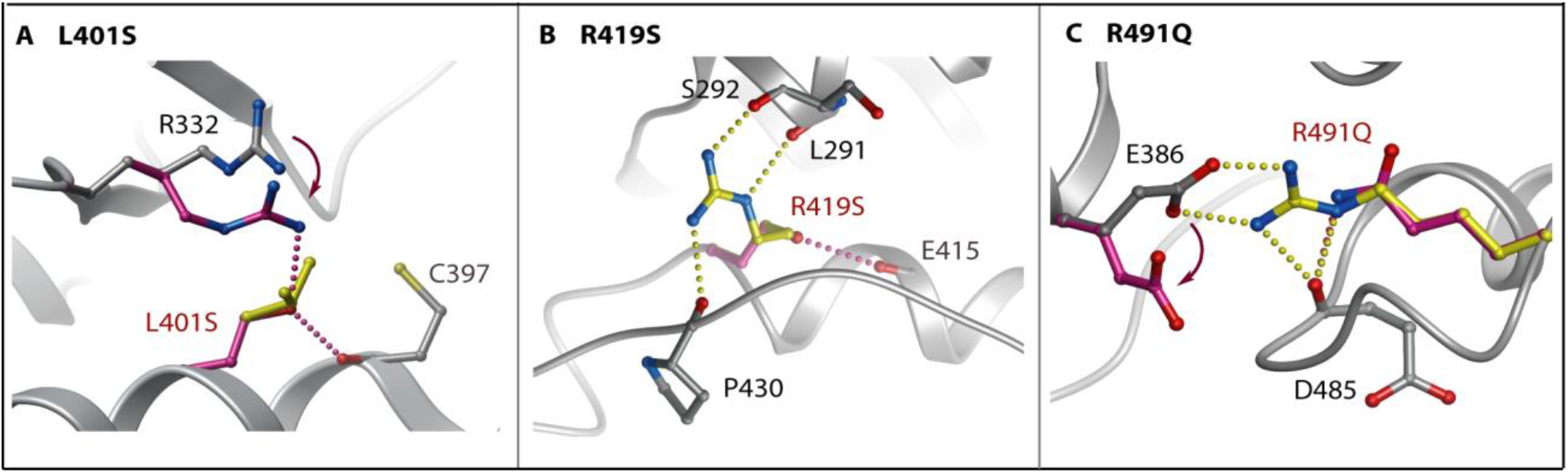
Modelling of PAH-associated mutations disrupting buried interactions. Structural models were calculated using the Eris server. Changes in the mutant structure are coloured purple and overlaid onto the wild-type residue coloured yellow. Hydrogen bonds in the wild-type (yellow) and clashes in the mutant structure (purple) are shown by spheres and dashed lines, respectively.

**Figure 7.**
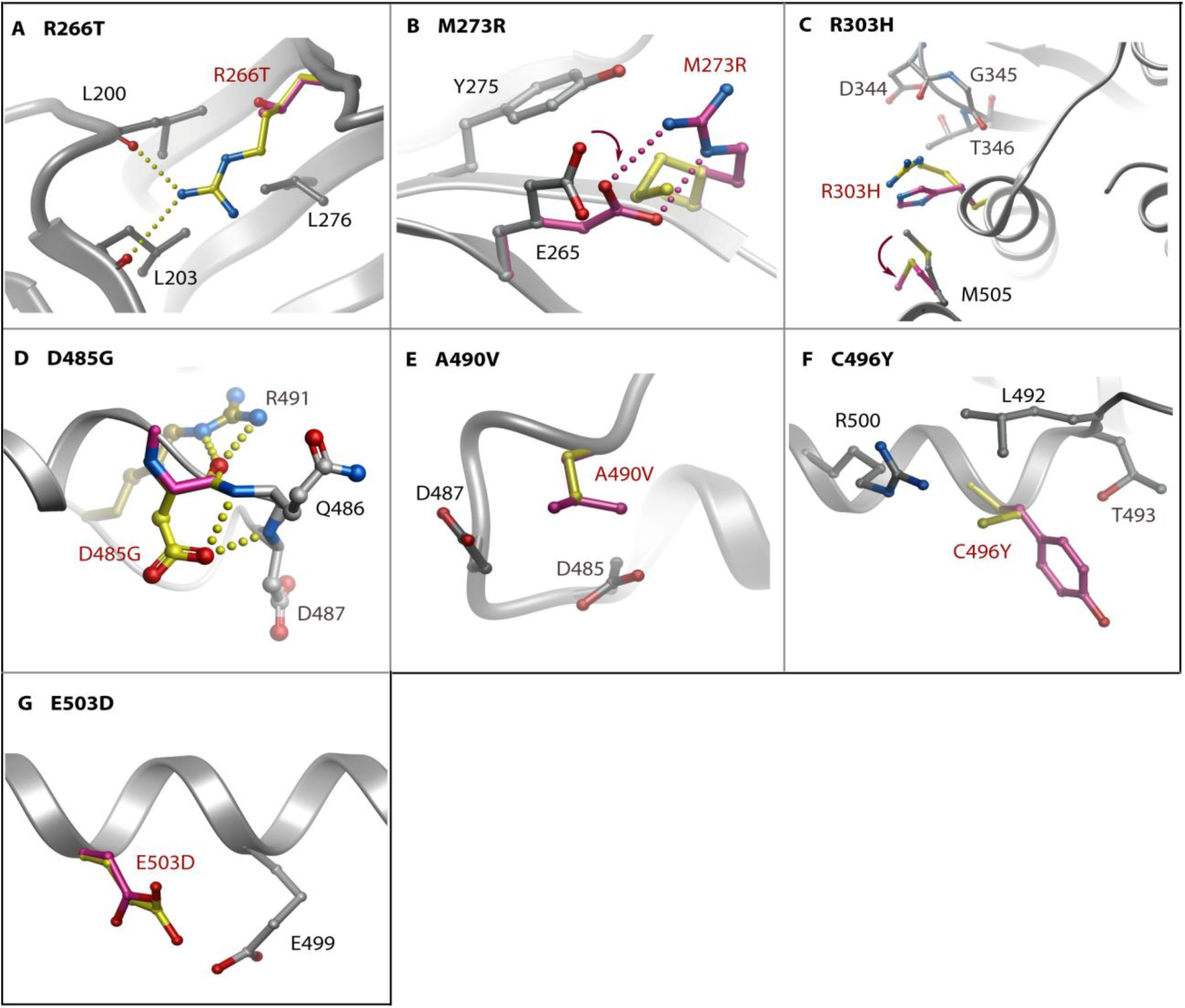
Modelling of PAH-associated mutations that are surface exposed in the BMPR2 structure. Structural models were calculated using the Eris server. Changes in the mutant structure are coloured purple and overlaid onto the wild-type residue coloured yellow. Hydrogen bonds in the wild-type (yellow) and clashes in the mutant structure (purple) are shown by spheres and dashed lines, respectively.

## Acknowledgements

The authors would like to thank the Swiss Light Source (SLS) station PX10 for beamtime. The SGC is a registered charity (number 1097737) that receives funds from AbbVie, Bayer Pharma AG, Boehringer Ingelheim, Canada Foundation for Innovation, Eshelman Institute for Innovation, Genome Canada, Innovative Medicines Initiative (EU/EFPIA) [ULTRA-DD grant no. 115766], Janssen, Merck KGaA Darmstadt Germany, MSD, Novartis Pharma AG, Ontario Ministry of Economic Development and Innovation, Pfizer, São Paulo Research Foundation-FAPESP, Takeda, and Wellcome [106169/ZZ14/Z].

## Author Contributions

A.N.B. and F.v.D designed the research. C.T. performed protein expression, purification and crystallisation. A.C. solved the structure. A.C. and A.N.B prepared the figures and wrote the initial draft manuscript. All authors approved the final manuscript.

